# The reovirus σ3 protein induces proteasomal degradation of IKKβ to modulate the host response to infection

**DOI:** 10.64898/2026.01.08.698360

**Authors:** Claudia Antonika, Durbadal Ojha, Pranav Danthi

**Author notes:** To whom correspondence should be addressed: Department of Biology, Indiana University, Bloomington, IN 47405.

## Abstract

Reovirus strains differ in their capacity to modulate host antiviral signaling, but the viral determinants underlying these differences have remained unclear. In this study, we identify the outer capsid protein σ3 as the viral determinant responsible for inducing the loss of the NF-κB kinase, IKKβ. We demonstrate that introduction of the σ3 encoding gene segment of strain T3A in the background of strain T3D^L^ recapitulates IKK loss observed during T3A infection. Our results indicate that *de novo* expression of T3A σ3 during infection is necessary to mediate the loss of IKK. Further, the presence of T3A σ3 alone is sufficient. We show that T3A σ3 interacts with the IKK complex and targets IKKβ for degradation through a Cullin-RING ligase–proteasome dependent mechanism. Loss of IKKβ results in impaired NF-κB activation and reduced IFN-β expression. Consequently, the T3D^L^/T3A S4 monoreassortant exhibits a replication advantage compared to T3D^L^. Collectively, these findings uncover a previously uncharacterized mechanism by which reovirus σ3 suppresses innate antiviral signaling through targeted degradation of IKKβ, providing new insight into reovirus mediated immune modulation.

**IMPORTANCE:** Successful viral infection and replication requires that the virus suppress host immune defenses, particularly innate immunity, which relies on signaling cascades to produce antiviral cytokines. Many viruses encode proteins to target key signaling molecules, such as those in the NF-κB pathway, but the exact mechanisms of evasion vary across viral families and even between closely related strains. In this study, we uncover a novel function of the reovirus outer capsid protein σ3 to promote IKKβ degradation. Our findings reveal that σ3 associates with a complex of proteins that includes IKKβ, a central kinase in the NF-κB pathway, to mediate its degradation via the proteasomal pathway. σ3 mediated IKKβ loss decreases type I IFN expression during infection and enhances viral replication. Coupled with previous observations that σ3 from other reovirus strains antagonizes NF-κB by altering its transcriptional activity, this work demonstrates that this conserved viral protein adopts strain-specific strategies to subvert host defenses.

## INTRODUCTION

The innate immune system provides the first line of defense against viral infections. A central component of this defense is the type I interferon (IFN) response, which plays a pivotal role in restricting viral replication and spread. For RNA viruses, the detection of viral RNA is mediated by endosomal sensors, such as Toll-like receptors (TLRs), and cytoplasmic sensors, RIG-I-like receptors (RLRs)(1). These pattern recognition receptors activate signaling cascades that converge on transcription factors such as Nuclear Factor κB (NF-κB) and Interferon Regulatory Factor 3 (IRF3), leading to the expression of type I IFNs and interferon-stimulated genes (ISGs). ISGs, in turn, establish an antiviral state in both infected and neighboring cells (2). Given the efficacy of IFN responses in restricting infections, viruses have evolved a range of mechanisms to subvert these responses. Many of these strategies involve sequestering, degrading, or inactivating key components of the interferon (IFN) signaling pathway. For instance, RSV ns2 protein sequesters RIG-I to limit nucleic acid sensing (3), rotavirus VP3 induces the degradation of the signaling adaptor MAVS (4), SARS-CoV-2 nsp6 and nsp13 inactivate the kinase TBK1 and the transcription factor IRF3 (5).

A key regulator within the antiviral signaling network is inhibitor of κB (IκB) kinase beta (IKKβ), the catalytic subunit of the IKK complex, which also includes IKKα and the regulatory subunit IKKγ/NEMO(6). Upon activation, the IKK complex phosphorylates IκBα, targeting it for proteasomal degradation. This degradation liberates NF-κB, allowing it to translocate to the nucleus and initiate transcription of a wide range of genes involved in inflammation, antiviral immunity, and cell survival (6).

Mammalian orthoreovirus (reovirus), a double-stranded RNA (dsRNA) virus, triggers IFN production via RIG-I-like receptor (RLR)-mediated sensing of its genomic RNA (7). The activation of NF-κB is essential for the induction of type I IFNs during reovirus infection. Accordingly, reovirus has evolved strategies to subvert the activation of NF-κB and limit IFN expression (8, 9). Previous work demonstrates that during infection with some reovirus strains, the transcriptional activity of NF-κB is blocked without affecting the levels or activity of upstream signaling components (10). In contrast, infection with the T3A strain of reovirus results in the loss of IKKβ, thereby suppressing NF-κB nuclear translocation(8). However, the viral factor responsible for IKKβ degradation by T3A and the underlying mechanism remained unclear.

In this study, we employed a combination of reverse genetics, viral reassortants, and ectopic expression systems to uncover the mechanism by which T3A infection results in loss of IKKβ. We identify that outer capsid protein σ3 from the T3A strain (T3A σ3) is both necessary and sufficient for inducing IKKβ loss. We demonstrate that T3A σ3 associates with the IKK complex and mediates IKKβ degradation via the Cullin ubiquitin ligase and proteasome dependent pathway. This targeted loss of IKKβ impairs type I IFN induction and promotes viral replication. Together, our findings establish T3A σ3 as a viral immune antagonist that degrades IKKβ, thereby impairing host antiviral defenses. Coupled with previous work, the findings presented here highlight the strain-specific functional diversity of reovirus σ3 proteins in their capacity to block NF-κB and underscores the critical role of NF-κB signaling in shaping the outcome of reovirus infection.

## RESULTS

### IKKβ loss following reovirus infection is influenced by the σ3 encoding S4 gene

Previous studies have demonstrated that infection with the T3A strain of reovirus leads to a decrease in levels of IKKβ (8). However, the specific viral factors or the underlying mechanism responsible for this effect have not been identified. During analysis of a monoreassortant virus, T3D^L^/T3A S4, in which the S4 gene segment encoding σ3 from the T3A strain was introduced into the T3D^L^ background, we found that infection led to a marked reduction in IKKβ protein levels, comparable to that observed with the T3A strain (Figure 1A). In contrast, infection with T3D^L^ had no impact on IKKβ expression. Parallel analysis of viral gene expression indicated that loss of IKKβ did not correlate with viral gene expression. Consistent with this, we found that under conditions of the experiments performed above, T3D^L^ and T3D^L^/T3A S4 replicated to comparable levels (data not shown). These findings suggest that strain dependent differences in IKKβ loss between T3D^L^ and T3A are impacted by the σ3 encoding S4 gene segment. To determine whether the reduction in IKKβ levels was due to repression in the transcription of the IKKβ gene, we measured IKKβ mRNA levels in cells infected with T3D^L^ and T3D^L^/T3A S4. RT-qPCR analysis at 24 h following infection revealed no significant differences in IKKβ mRNA levels between cells infected with the two virus strains (Figure1C). These data indicate that reduction in IKKβ occurs post-transcriptionally.

**FIG 1.**
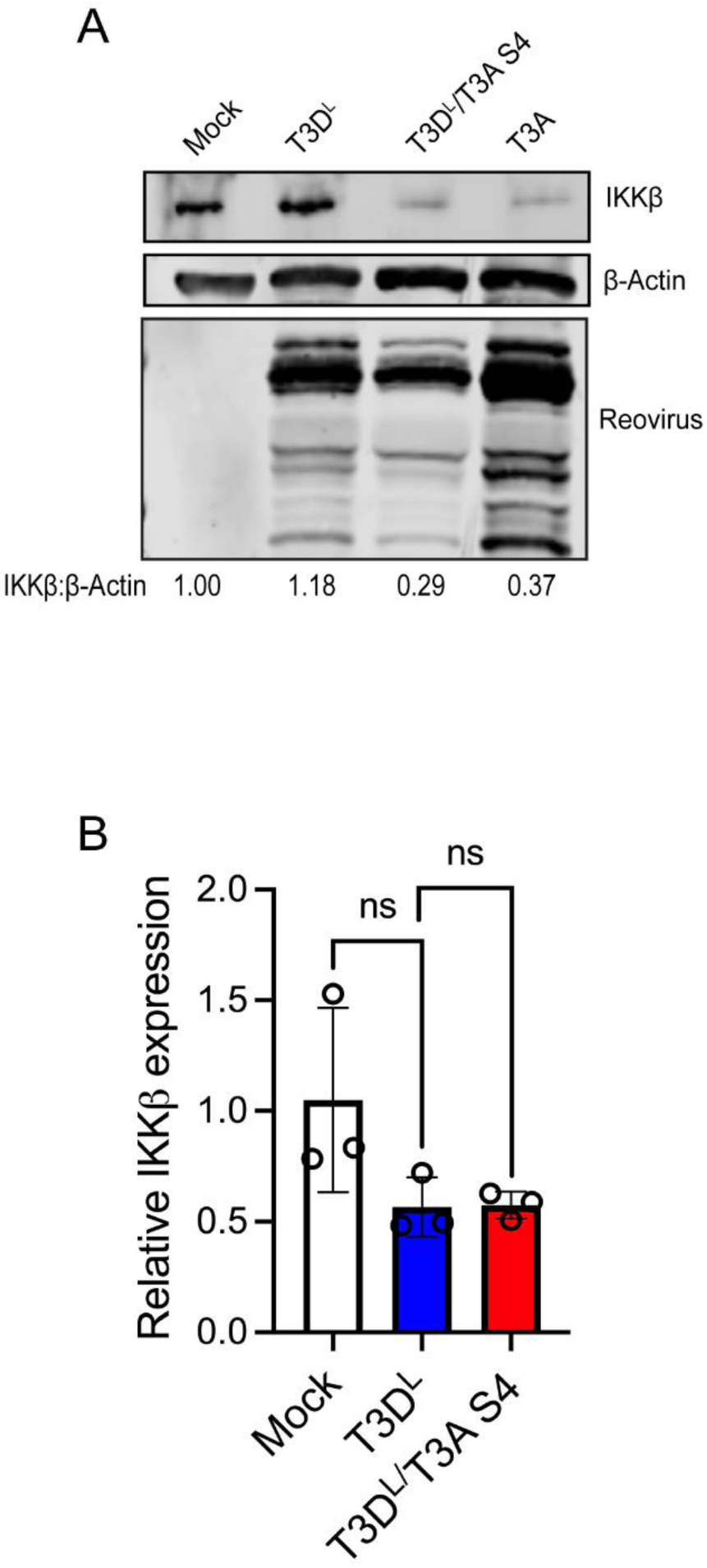
The T3A S4 gene post transcriptionally alters IKKβ levels in infected cells. ATCC L929 were adsorbed with PBS (Mock), T3D^L^, T3D^L^/T3A S4, or T3A at multiplicity of infection (MOI) 10 PFU/cell. Following incubation at 37°C for 20 h, the cells were harvested. (A) Whole cells lysate from infected cells were immunoblotted using α-reovirus, α-IKKβ, and α-β-actin antibody. (B) RNA was extracted from cells and the levels of IKKβ mRNA relative to GAPDH mRNA was measured using RT-qPCR and comparative C_T_ analysis. The ratio of each mRNA relative to GAPDH in cell treated with PBS (Mock) was set to 1. The mean values and SD are shown. **, *P* values were determined by One-Way ANOVA with Dunnet’s multiple correction test. ns, not significant, * *P<0.05, \*\**.

### Newly synthesized T3A σ3 mediates IKKβ loss

σ3 is a multifunctional protein (11, 12). As part of the viral outer capsid, it controls cell entry of reovirus. During viral entry, σ3 present in incoming particles is degraded by host proteases(13). Therefore, it is unlikely that the σ3 from incoming virions could directly impact IKKβ levels. Yet, it remains possible that virion associated σ3 alters virus entry kinetics or efficiency and indirectly influences IKKβ levels. To rule out the role of incoming σ3 in IKKβ loss, we generated infectious subviral particles (ISVPs) from virions by *in vitro* chymotrypsin treatment. This treatment removes σ3 from the particle surface. Because only the σ3 encoding S4 gene is different between T3D^L^ and T3D^L^/T3A-S4, ISVPs generated from virions of these two strains are identical. When infection was initiated with ISVPs derived from these strains, we found that whereas infection with T3D^L^ ISVP failed to induce IKKβ loss, infection with ISVPs of the T3D^L^/T3A S4 monoreassortant still led to IKKβ degradation (Figure 2A). These results indicate that capsid associated σ3 is not required for mediating IKKβ loss.

**FIG 2.**
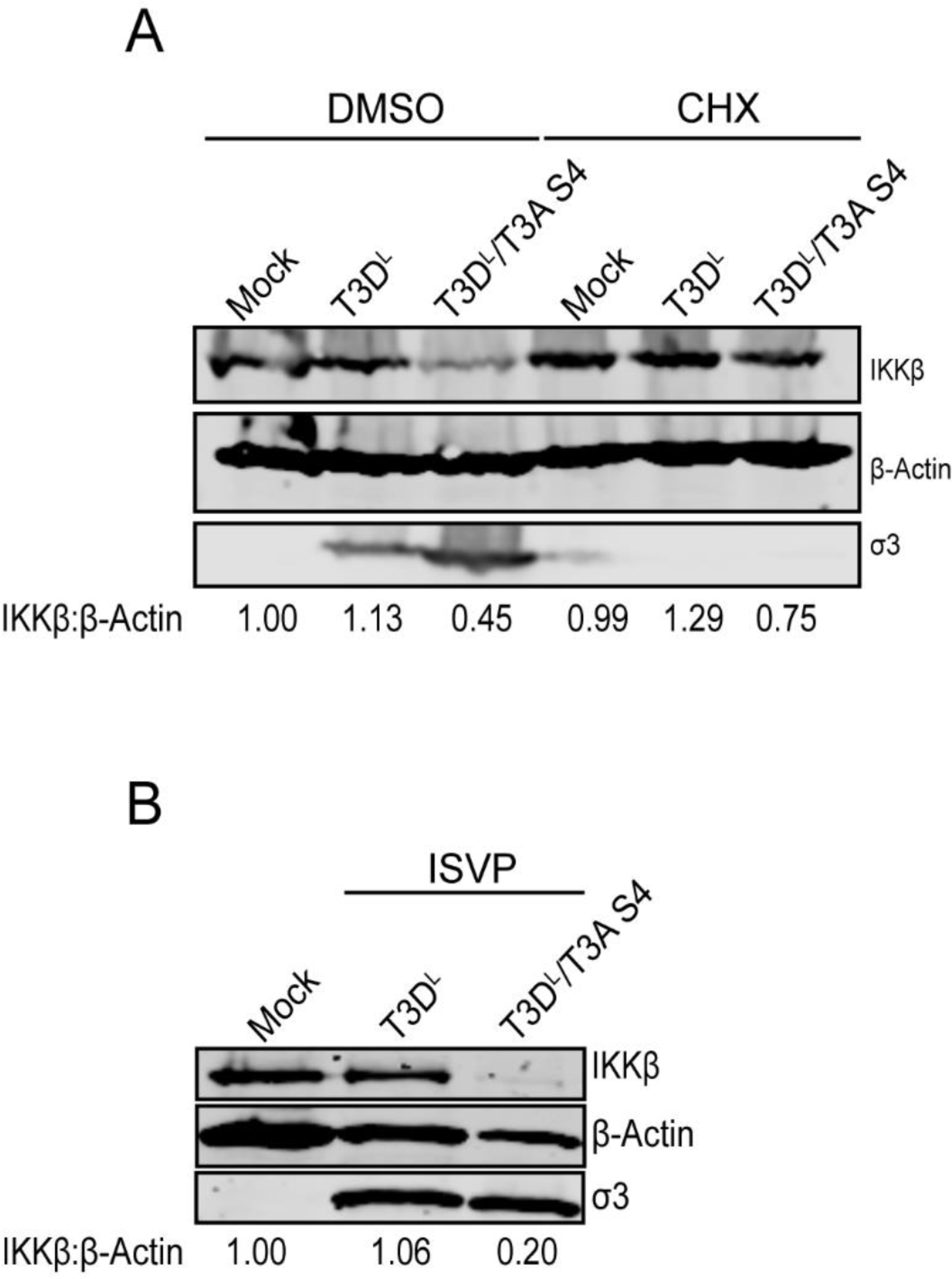
Virion associated σ3 does not contribute to IKKβ loss. (A) ATCC L929 were adsorbed with PBS (Mock) or ISVPs of T3D^L^ and T3D^L^/ T3A S4 at MOI 10 PFU/cell for 1 h at room temperature. At 24 h following infection, the cells were harvested and immunoblotted using α-σ3, α-IKKβ, and α-β-actin antibodies. (B) ATCC L929 were adsorbed with PBS (Mock) or virions of T3D^L^ and T3D^L^/ T3A S4 at MOI 10 PFU/cell for 1 h at room temperature. After adsorption, the cells were treated with 10 μg/mL cycloheximide and incubated for 20 h The cells were harvested and subjected for immunoblotting using α-σ3, α-IKKβ, and α-β-actin antibody.

Later in infection, σ3 is produced *de novo* following transcription of the viral genome. Newly synthesized σ3 plays additional roles in viral replication and in controlling host responses. To determine whether IKKβ loss observed following infection is mediated by newly synthesized σ3, we determined whether the reduction in IKKβ levels required the synthesis of new viral proteins. For these experiments, we treated infected cells with cycloheximide (CHX) to block protein synthesis and then assessed IKKβ levels by immunoblotting (Figure 2B). We found that blocking of viral protein synthesis (as measured by accumulation of σ3 in infected cells), prevented IKKβ degradation during T3D^L^/T3A S4 infection. These data suggest that *de novo* synthesis of viral proteins is required to induce IKKβ loss.

### T3A σ3 expression is necessary and sufficient for reduction of IKKβ levels

Given the effect of newly synthesized viral proteins on IKKβ levels, we asked if expression of σ3 during infection was required for this effect. For these experiments, we used siRNA to knock down σ3 transcripts in infected cells. Because no loss of IKKβ is observed following T3D^L^ infection, knockdown of T3D^L^ σ3 did not impact IKKβ levels. In contrast, upon knockdown of T3A σ3, we observed that T3D^L^/T3A S4 induced IKKβ loss was partially restored (Figure 3A). These data indicate that T3A σ3 expression is necessary for IKKβ degradation.

**FIG 3.**
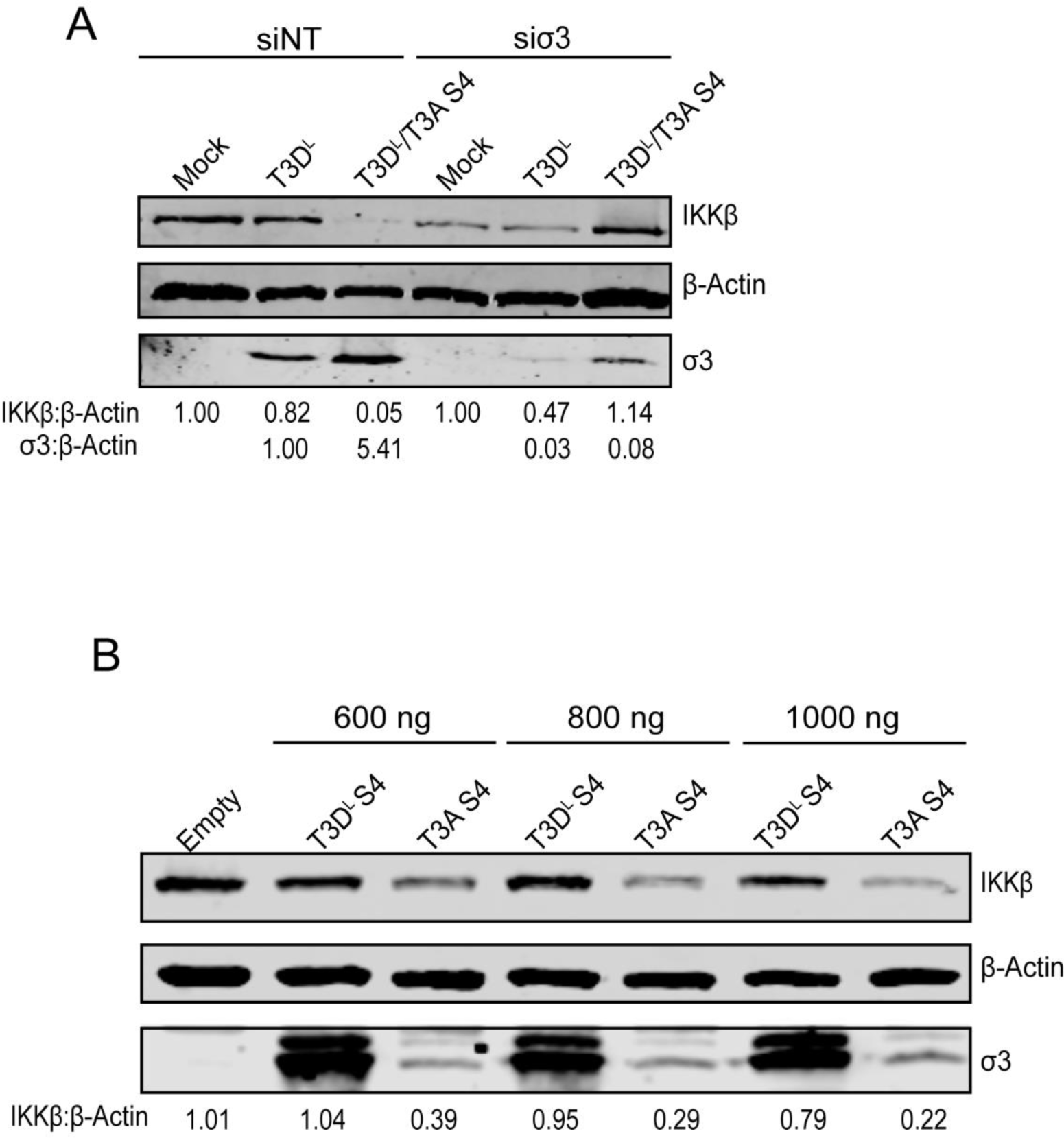
Expression of T3A σ3 is necessary and sufficient for changing IKKβ levels. (A) L929 cells were transfected with either β-gal siRNA for the non-targeting (NT) or σ3 siRNA using INTERFERin. At 24 h following transfection, the cells were infected with PBS (Mock), T3D^L^ or T3D^L^/T3A S4 at MOI of 5 PFU/cell and incubated for an additional 24 h. (B) HEK293T were transfected with the indicated concentration of empty vector or vector expressing T3D^L^ σ3 or T3A σ3. The cells were harvested at 48 h following transfection. (A,B) The cells were harvested and immunoblotted using α-σ3, α-IKKβ, and α-β-actin antibodies.

It is possible that σ3 knockdown prevents IKKβ degradation because T3A σ3 directly mediates this effect. However, because σ3 knockdown can have other effects on infection, it is possible that the effect of σ3 knockdown on IKKβ is indirect. To distinguish between these possibilities, we next tested whether σ3 expression alone is sufficient to induce this effect. For this experiment, increasing amounts of T3D^L^ or T3A σ3 expression vectors were transfected into cells. Despite the fact that the same vector backbone was used for each, at each plasmid concentration, T3D^L^ σ3 accumulated to higher levels than T3A σ3. Nonetheless, we observed that only T3A σ3 mediated dose dependent IKKβ loss (Figure 3B). These data indicate that expression of T3A σ3 alone is sufficient to induce IKKβ loss. Further, our results highlight that the capacity of σ3 to induce IKKβ loss varies with functional differences between σ3 from two different strains and does not correlate with the amount of each type of σ3 present in the cell.

### T3A σ3 targets IKKβ for Degradation via the Cullin-RING Ligase (CRL)-proteasomal pathway

Degradation of cytoplasmic proteins in mammalian cells primarily occurs through two pathways (14). Proteins may be subject to autophagosome mediated degradation requiring lysosomal enzymes. Alternatively, proteins may be degraded via the ubiquitin-proteasomal system. Our previous work showed that T3A-induced IKKβ loss is not rescued by ammonium chloride, an inhibitor of lysosomal acidification (8). These results raised the possibility that the ubiquitin-proteasomal pathway is involved in the loss of IKKβ. However, because proteasome inhibitors block efficient entry of reovirus into cells (15), the role of this pathway in diminishment of IKKβ levels following infection has not been formally tested.

Our observation that IKKβ levels in cells transfected with a vector encoding T3A σ3 is diminished allowed us to evaluate the contribution of proteasomal degradation in σ3-mediated IKKβ turnover. For these experiments, cells transfected with T3A σ3 expression vector were treated with a commonly used proteasome inhibitor, MG132, and IKKβ levels were analyzed by immunoblotting. We found that MG132 treatment prevented IKKβ degradation observed in T3A σ3 expressing cells that are treated with DMSO. MG132 treatment of T3A σ3 expressing cells brought IKKβ levels to those seen in cells transfected with empty vector (Figure 4A). Thus, our data indicate that IKKβ is turned over by the proteasome when T3A σ3 is expressed.

**FIG 4.**
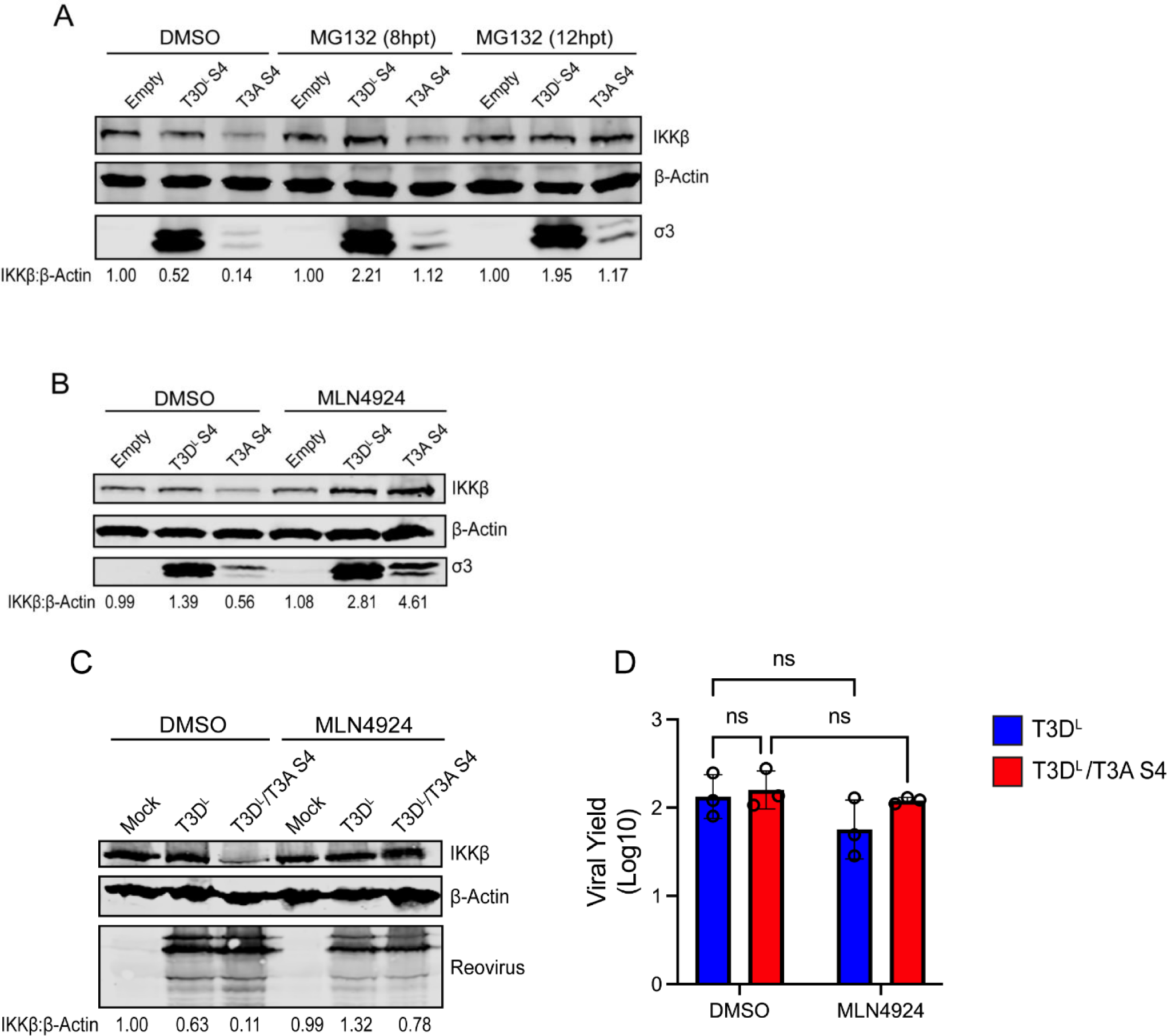
IKKβ loss requires the function of the proteasome and Cullin ubiquitin ligases. (A, B) HEK293T were transfected with 1.2 μg of empty vector or vector expressing T3DL σ3 or T3A σ3. 36 h post-transfection, cells were treated with DMSO or 25 μM of MG132 for 8 or 12 h (A) and 2 μM of MLN4924 for 20 h. (C and D) ATCC L929 were adsorbed with PBS, T3D^L^, T3D^L^/T3A S4 at MOI 10 PFU/cell for 1 h at room temperature. Following adsorption, the cells were treated with 2 μM MLN4924 for 20 h. (A,B,C) The cells were harvested and immunoblotted using α-σ3, α-reovirus, α-IKKβ, and α-β-actin antibody. (D) At 24 h, the virus was harvested by freeze-thaw and the titer was measured by plaque assay on L929 cells.

Ubiquitin-dependent proteasomal degradation requires a cascade of enzymes including E1 ubiquitin-activating enzymes, E2 conjugating enzymes, and E3 ubiquitin ligases. Among these, E3 ligases provide substrate specificity (16). We next sought to identify the class of E3 ligases involved in this process. E3 ligases fall into three main families, Really Interesting New Gene (RING), Homologous to the E6-AP Carboxyl Terminus (HECT), and RING-Between-RING (RBR), and among them, the Cullin-RING ligases (CRLs) represent the largest and most functionally diverse subclass(18). We therefore used MLN4924, a small molecule inhibitor of NEDD8-activating enzyme that specifically disrupts CRL function, to evaluate the contribution of CRLs to σ3-induced IKKβ loss. We treated control and σ3 expressing cells with MLN4924. MLN4924 treatment effectively prevented σ3-induced IKKβ degradation (Figure 4B), indicating that a CRL-type E3 ligase mediates this process.

Next, we assessed whether CRL activity is also required for IKKβ loss in the context of viral infection. MLN4924 treatment did not significantly inhibit viral protein expression in infected cells or affect viral replication (Figure 4D, E). Yet, MLN4924 blocked IKKβ degradation in cells infected with T3D^L^/T3A S4. Our results therefore indicate that CRL activity is necessary for σ3-mediated IKKβ degradation during infection (Figure 4C). Collectively, these findings suggest that IKKβ loss in transfected and infected cells occurs via the same mechanism and requires degradation via the CRL-proteasomal pathway.

### T3A σ3 forms a complex with IKKβ

Many viruses promote host protein degradation by using viral proteins to bind their target and bring in an E3 ligase (16). Based on this model, we hypothesized that T3A σ3 associates with IKKβ to recruit the ligase that mediates its degradation. To test this idea, we transfected cells with expression vectors for T3D^L^ or T3A σ3 along with a vector expressing FLAG-tagged IKKβ. To observe a potential interaction, the cells were treated with MLN4924 to prevent IKKβ degradation. Consistent with our earlier observations, T3A σ3 was expressed at lower levels than T3D^L^ σ3 in whole-cell lysates when lysates from an equivalent number of cells were resolved (Figure 5A). To enable comparison of the interaction capacity of T3D^L^ and T3A σ3 proteins, we used five times more lysate from T3A σ3 transfected cells for our immunoprecipitation compared to the remaining samples (Figure 5A, bottom panel). This adjustment brought the levels of T3D^L^ and T3A σ3 in a comparable range. Upon immunoprecipitation of IKKβ using anti-FLAG antibody, we found that both T3D^L^ σ3 and T3A σ3 were detected. However, T3A σ3 consistently showed stronger binding to the IKKβ containing complex compared to T3D^L^ σ3 (Figure 5A), suggesting there may be strain-specific differences in binding affinity or interaction stability. Reovirus σNS, a similarly sized viral protein, did not demonstrate any interaction with the IKKβ complex. To further validate this interaction, we transfected cells with empty or T3A σ3 expression vector along with an expression vector for Flag tagged IKKβ. We found that upon immunoprecipitation of T3A σ3 using an anti-σ3 antibody, IKKβ was pulled down (Figure 5B). These results support a model in which σ3 associates with IKKβ directly or with a complex that also contains IKKβ. For the T3A strain, this interaction facilitates targeting of IKKβ to the CRL-proteasomal degradation pathway.

**FIG 5.**
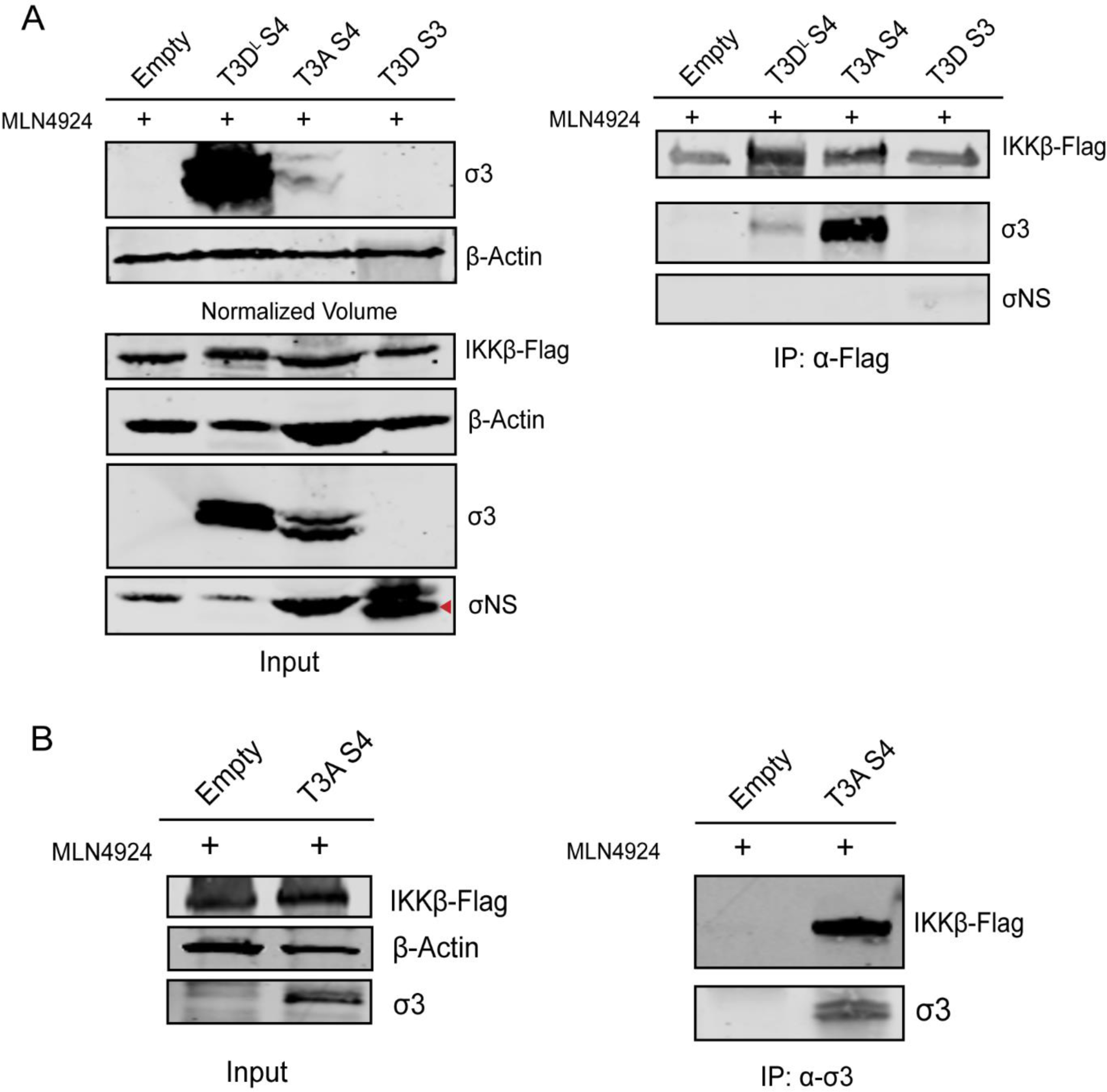
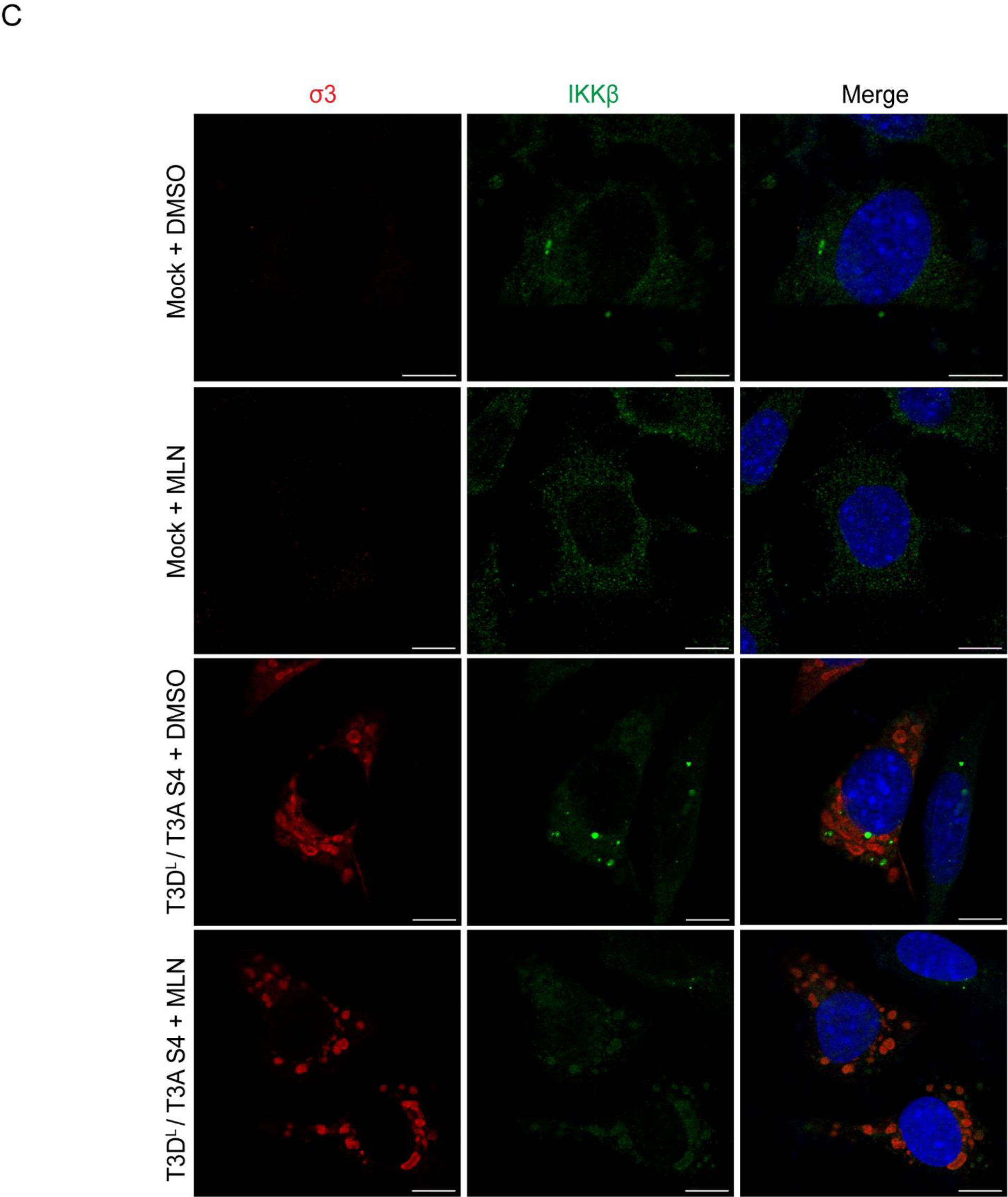
T3A σ3 associates with IKKβ. (A,B) HEK293T were transfected with 2 μg of IKKβ-Flag expression vector and either 3 μg of empty vector or vector expressing T3D^L^-σ3, T3A-σ3, or σNS. Following incubation at 37°C for 24 h, the cells were treated with 2 μM MLN4924 and incubated for another 20 h. The cells were harvested and subjected to immunoprecipitation with α-flag (A) or α-σ3 (B) antibody. The input and the IP lysates were immunoblotted with α-σ3, α-flag, σ-β-actin, or α-σNS antibody. (C) L929 were infected with T3D^L^/T3A S4 at MOI 10. After adsorption, the cells were treated with MLN4924 or DMSO at dose of 2μM. At 24h, cells were immunolabeled with σ3 (red), IKKβ (green) and host cell nuclei (blue) and finally imaged with a confocal microscopy. Images were deconvolved using Huygens Essential (17.10) software and scale bar (10 µm) were added using ImageJ software. Images are representative of three independent experiments.

We found it difficult to detect the association of endogenous IKKβ with σ3 in infected cells, likely due to a lower-level expression of endogenous IKKβ, the quality of commercially available anti-IKKβ antibodies or both. Therefore, to further assess the association between T3A σ3 and endogenous IKKβ in infected cells, we used immunofluorescence microscopy. Endogenous IKKβ exhibited a diffuse cytoplasmic distribution in uninfected cells treated with either DMSO or MLN4924. In contrast, in cells infected with T3D^L^/T3A S4 and treated with DMSO, IKKβ fluorescence intensity was reduced, likely due to degradation of endogenous IKKβ. In infected cells treated with MLN4924, IKKβ signal intensity was higher and appeared to co-localize with T3A σ3 (Figure 5C). These results indicate that T3A σ3 associates with endogenous IKKβ in infected cells and that this association is more readily detected when CRL-dependent degradation is inhibited.

### The presence of μ1 weakens T3A σ3 interaction with IKKβ

During infection, σ3 can exist in distinct forms: freely in the cytoplasm or nucleus, or in complex with its viral partner outer capsid protein, μ1, which is associated with lipid droplets and viral inclusions (17). Earlier studies demonstrate that σ3 and μ1 influence one another’s functions when they interact (18, 19). In particular, σ3 binds dsRNA only in its unassociated form (19). We therefore hypothesized that the form of σ3 not bound to μ1 is responsible for mediating IKKβ loss. To test this idea, we co-transfected cells with plasmids expressing T3A σ3 and μ1 and assessed IKKβ levels. Co-expression of μ1 inhibited the capacity of σ3 to induce IKKβ loss (Figure 6A). These data indicate that μ1-bound σ3 is functionally sequestered and unable to promote IKKβ loss. Based on this result, we predicted that the presence of μ1 would prevent the association of σ3 to IKKβ. Indeed, consistent with data presented above, T3A σ3 was found in association with IKKβ. This association was dramatically reduced when μ1 was also expressed in the cell (Figure 6B, right panel). These findings suggest that only μ1-free σ3 can interact with IKKβ.

**FIG 6.**
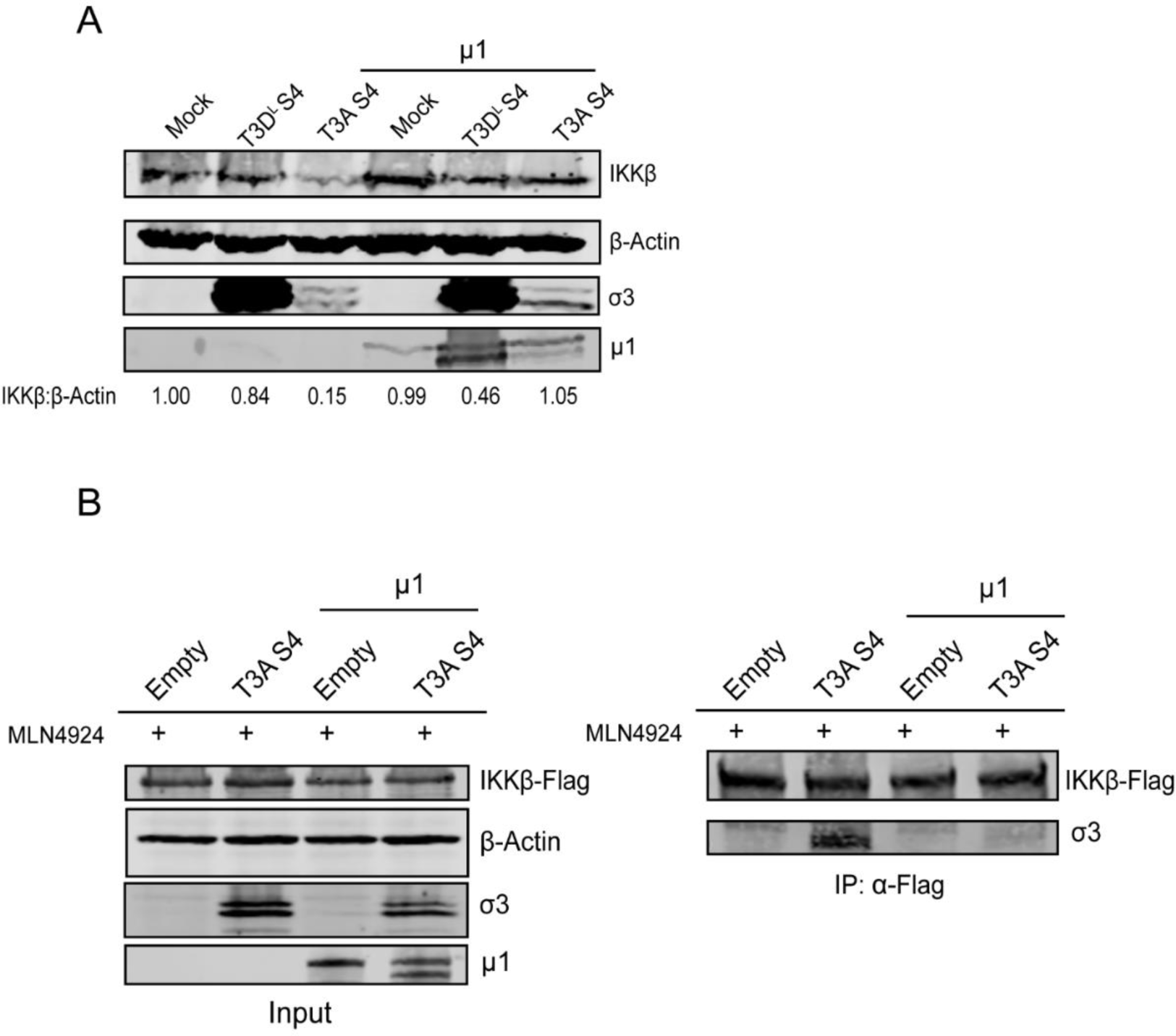
The presence of μ1 prevents IKKβ turnover by T3A-σ3. (A) HEK293T were transfected with 1.2 μg of empty vector or vector expressing T3DL σ3 or T3A σ3 along with 0.8 μg of expression vector for T3D^F^-μ1. 48 h after transfection, the cells were harvested and immunoblotted using α-σ3, α-flag, and α-β-actin antibodies (B) HEK293T were transfected with transfected with 3 μg of empty vector or vector expressing T3DL σ3 or T3A σ3 along with 2μg of IKK-β flag expression vector and 1 μg of T3D-μ1 expression vector. Following incubation at 37°C for 24 h, the cells were treated with 2 μM MLN4924 and incubated for an additional 20h. The cells were harvested and subjected to immunoprecipitation with α-flag antibody to precipitate IKKβ-flag. (A,B) The cells lysates were harvested and immunoblotted using α-σ3, α-flag, and α-β-actin, or α-reovirus antibodies.

### T3A σ3 suppresses expression of NF-κB target genes

The genome of reovirus is recognized by RIG-I family cytoplasmic sensors to induce the activation of NF-κB. Since IKKβ is a core component of the IKK complex required for NF-κB activation, we sought to determine if loss of IKKβ results in reduction in expression of NF-κB target genes. Using RT-qPCR, we found that expression of a known specific NF-κB target, A20, is significantly reduced in T3D^L^/T3A S4 infected cells (Figure 7A). The type 1 IFN, IFNβ is another NF-κB target. IFNβ expression is dependent on the activation of NF-κB and IRF3 pathways (20). However, since under some conditions, IFNβ can also be expressed without the function of NF-κB (21), we first confirmed using RT-qPCR that IFNβ induction during reovirus infection in our experimental set up is dependent on IKKβ and NF-κB. We found that inhibition of IKKβ significantly reduced IFNβ mRNA levels in reovirus infected cells (Figure 7B). Based on this, we hypothesized that infection with T3D^L^/T3A S4 would result in lower type I IFN expression due to its ability to degrade IKKβ. Consistent with this hypothesis, IFNβ mRNA expression was significantly reduced in cells infected with T3D^L^/T3A S4 compared to the T3D^L^ parental strain (Figure 7C). Because IFNβ induction following reovirus infection is affected by cell entry events and because σ3 properties can affect entry, it was important to ascertain whether the difference in IFNβ induction between T3D^L^ and T3D^L^/T3A S4 is related to the NFκB inhibitory effect of newly synthesized σ3. For this, we measured IFNβ expression following infection initiated by *in vitro* generated ISVPs of each of these reovirus strains. ISVP infection recapitulated the strain specific difference in IFNβ mRNA observed with virions, confirming that newly synthesized σ3 is responsible for suppressing type I IFN responses (Figure 7D). Together, these findings suggests that IKKβ loss mediated by T3A σ3 dampens innate immune responses by suppressing NF-κB-dependent type I IFN signaling.

**FIG 7.**
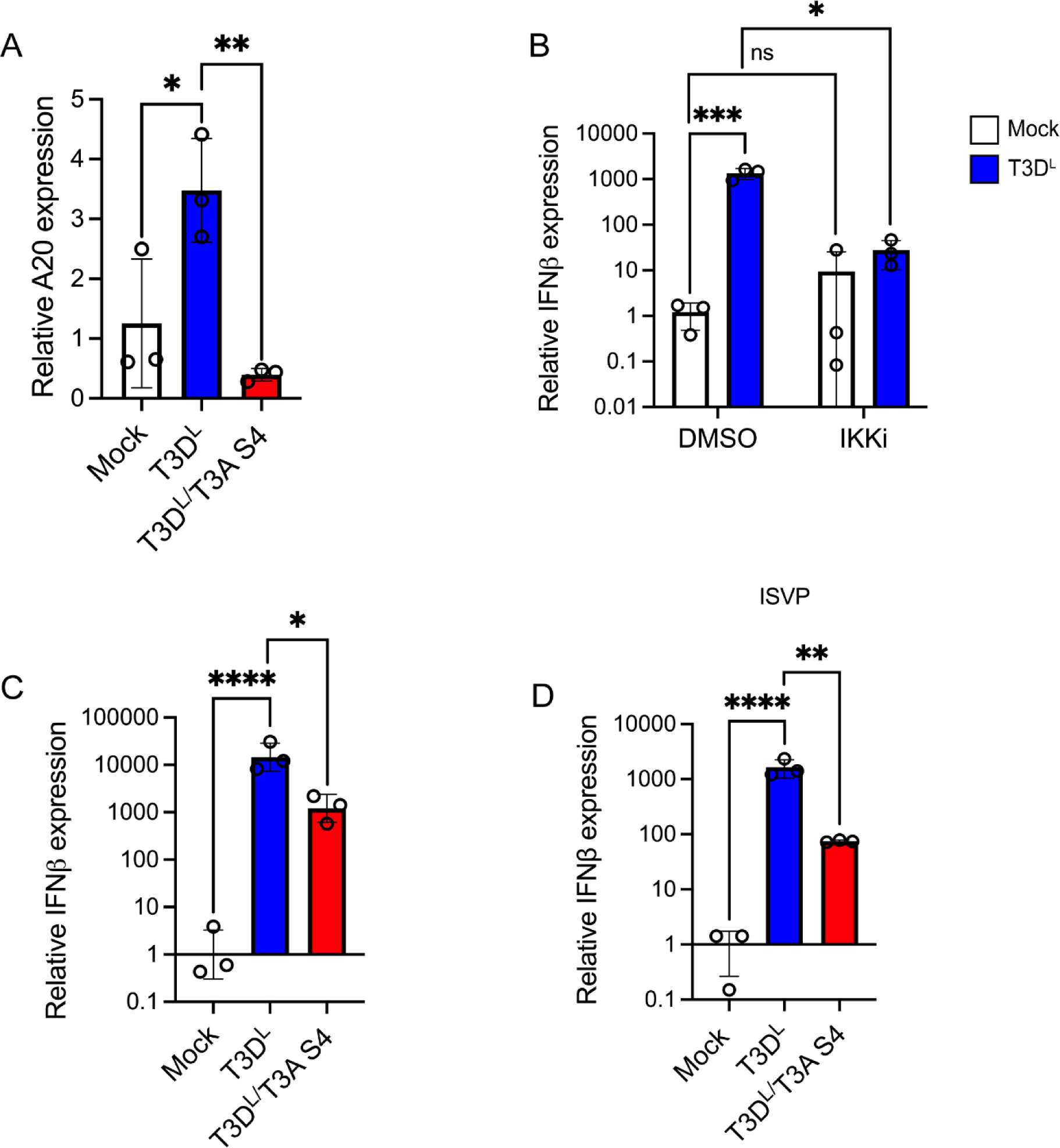
Expression of NFκB target genes is reduced following infection with T3D^L^/ T3A S4 (A-D) ATCC L929 were absorbed with PBS (mock), or 10 PFU/cell of T3D^L^, T3D^L^/T3A S4, T3D^L^ or T3D^L^/T3A S4 ISVPs. (B) After adsorption, the cells were treated with DMSO or 10 μM of IKK inhibitor (BAY 65-1942) at 37°C for 20 h. (A-D) RNA was extracted from the harvested cells and the levels of A20 and IFNβ mRNA relative to GAPDH mRNA was measured using RT-qPCR and comparative C_T_ analysis. The ratio of A20 or IFNβ mRNA relative to GAPDH in cell treated with PBS (Mock) was set to 1. The mean values and SD are shown. *P* values were determined by Two-Way ANOVA with Tukeys multiple comparisons test. ns, not significant, * *P<0.05, ** P<0.005, *** P<0.0005*.

### T3A σ3-driven IKKβ degradation influences viral replication

Because T3D^L^/T3A S4 produces less IFNβ than T3D^L^, we hypothesized that this difference might influence viral replication. In earlier experiments using a high MOI of 10 (data not shown), we observed no difference in replication kinetics between the two viruses, likely because all cells were infected simultaneously, masking IFN-dependent paracrine effects that are thought to be dominant during reovirus infection (22). To better capture potential differences in antiviral responses, we infected L929 cells at an MOI of 1. At lower MOI, the effect of IFN becomes more apparent because uninfected bystander cells resist or diminish viral infection in an IFN-mediated manner. Under lower MOI conditions, at 24 h post infection, T3D^L^/T3A S4 generated a modestly higher viral yield than T3D^L^, although this difference is diminished by 48 h (Figure 8). These findings align with the reduced IFN production caused by IKKβ degradation in T3D^L^/T3A S4, which allows more efficient viral replication.

**FIG 8.**
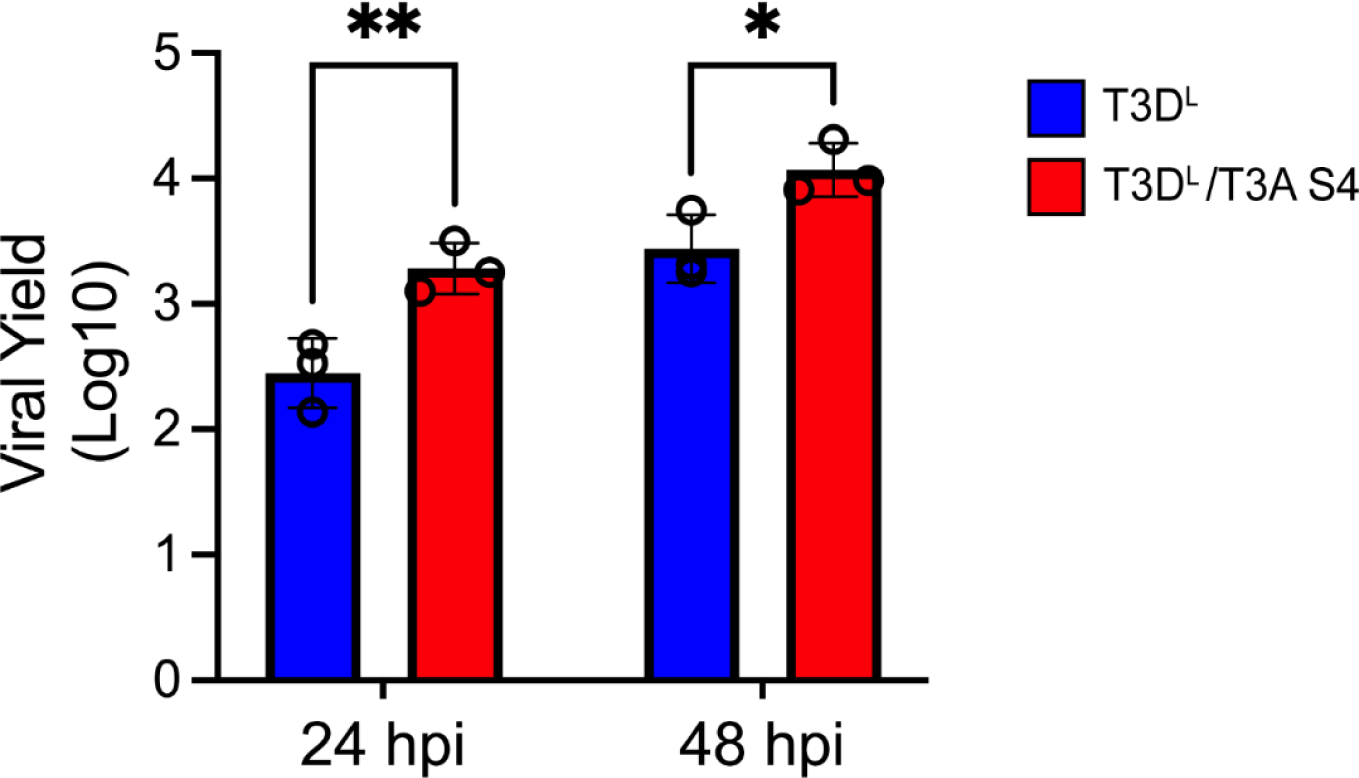
T3D^L^/ T3A S4 displays a replication advantage at lower MOI. ATCC L929 were adsorbed with PBS, T3D^L^ and T3D^L^/T3A S4 at multiplicity of infection (MOI) 1 PFU/cell. Following incubation at 37°C for the indicated time points, the virus was harvested by freeze-thaw and the titer was measured by plaque assay on L929 cells.

## DISCUSSION

Reovirus strain T3A suppresses NF-κB signaling by reducing IKKβ abundance (8), but the viral determinant responsible for this effect has remained unknown. Here, we identify σ3 from the T3A strain as the key factor that drives IKKβ degradation and clarify the mechanism by which this occurs. We show that newly synthesized σ3, rather than virion-delivered protein, is responsible for IKKβ loss during infection. Expression of T3A σ3 alone is sufficient to reduce IKKβ levels in a dose-dependent fashion, demonstrating that σ3 is both necessary and sufficient for this phenotype. IKK loss occurs post-transcriptionally and we further show that blocking CRL neddylation or proteasome function rescues IKKβ levels (Figure 9). Our results suggest that T3A σ3 mediated IKK turnover is due to the association of σ3 with a protein complex containing IKKβ. IKK degradation results in a lower-level induction of the innate immune response and consequently higher virus replication.

**Figure 9.**
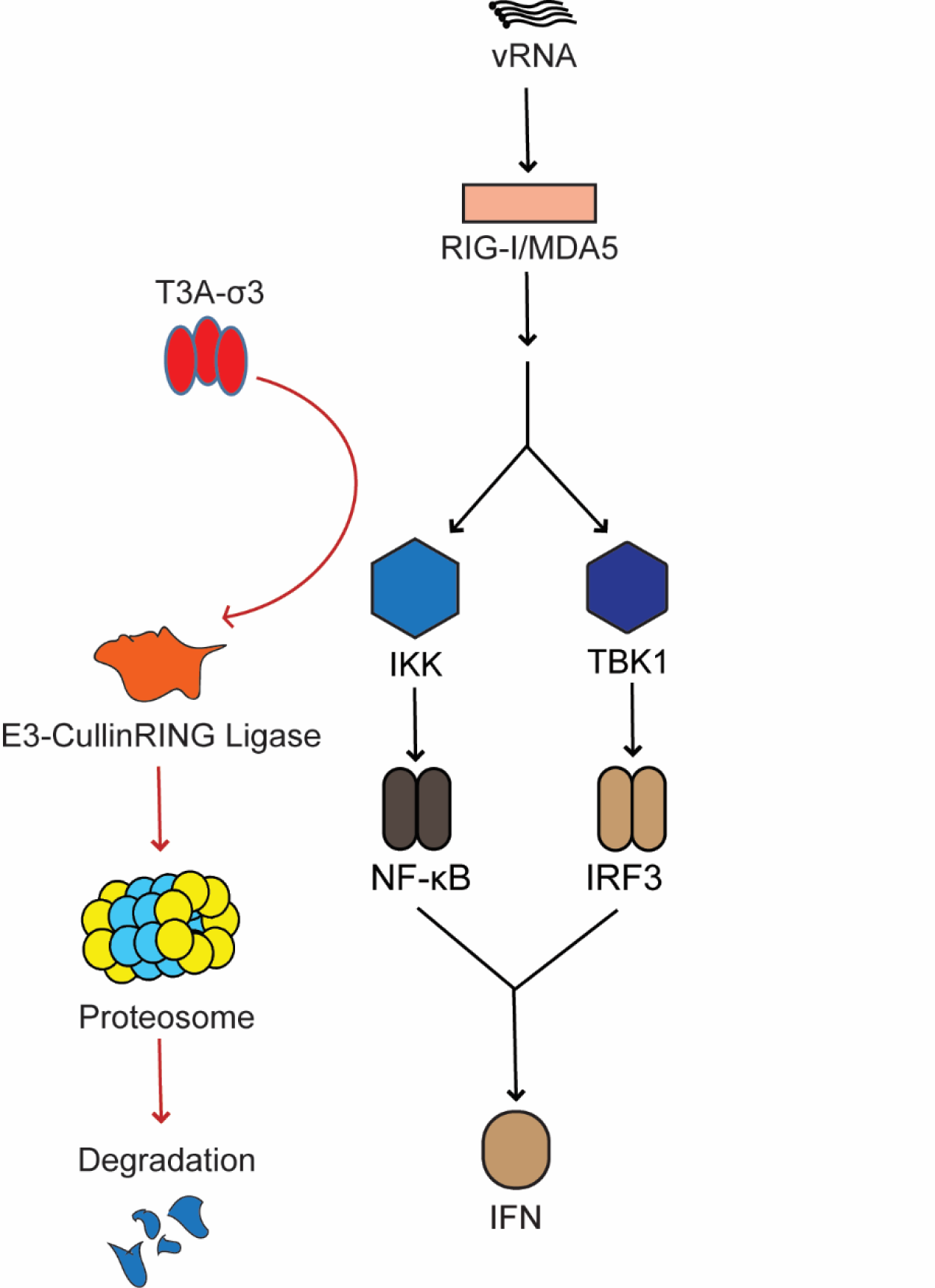
Model of T3A σ3 mediated IKKβ degradation. Newly synthesized σ3 from the T3A strain associates with an IKKβ and promotes CRL dependent, proteasome-mediated degradation of IKKβ. Loss of IKKβ dampens innate immune signaling and enhances viral replication.

The IKK kinase complex is a central regulator of NF-κB signaling, composed of two catalytic subunits (IKKα and IKKβ) and the regulatory subunit NEMO, along with multiple transient and stable accessory proteins. This assembly forms a large ∼700-900 kDa complex(6). In our study, we demonstrated that IKKβ is degraded by T3A σ3 and we also found that σ3 is associated with IKKβ. However, the mechanism by which σ3 induces IKKβ turnover remains unknown. While IKKβ was diminished by σ3, it is unclear whether σ3 is only targeting IKKβ or additional members of the complex. Our previous work found that NEMO was also degraded during T3A infection (8). This suggests that σ3 may likely induce degradation of additional members of the IKK complex. Which member(s) of the IKK complex are subject to ubiquitination and degradation currently remain unknown. Such experiments are further complicated by evidence that formation of the IKK complex depends on ubiquitin modification (23). Similarly, the direct binding partner of σ3 within this complex is unidentified. Given the number of accessory proteins within the complex, it is difficult to determine whether σ3 targets IKKβ directly or interacts with another protein that indirectly results in IKKβ precipitation in our assays.

Because inhibition of neddylation with MLN blocks IKKβ loss, we infer that one or more Cullin-RING ligase (CRL) is involved. CRL are modular complexes which consist of 7 different types of Cullin proteins serving as scaffolds. Each Cullin interacts with an RBX protein onto which E2 conjugated with a ubiquitin is loaded (16). Each Cullin uses a different adaptor protein that links the substrate receptor to the Cullin to put the substrate in close proximity to E2. As shown in figure 5, σ3 associates with IKKβ, this interaction with IKK complex may allow σ3 to recruit the Cullin. Though the type of Cullin needed for σ3 mediated IKKβ degradation remains to be identified, it is possible that σ3 may act as an adaptor protein to target IKKβ for degradation. Most Cullin adaptor proteins have specific substrates. This substrate may be post-translationally modified or contain a degron motif which is recognized by a specific CRL adaptor protein. For instance, the KEAP1 adaptor protein recognizes (D/N)XE(T/S)GE motif found in human IKKβ and promotes its ubiquitination by the Cullin 3 (CUL3)/ Ring-Box 1 (RBX1) dependent E3 ubiquitin ligase complex(24). This motif is lacking in murine IKKβ. Since our data suggest that that T3A σ3 mediates IKKβ degradation in both mouse and human cells, we expect that there exists another mechanism that is shared between mouse and human cells that mediates IKKβ turnover in the presence of σ3. For example, it is possible that σ3 may not directly recruit the Cullin scaffold but instead recruit other adaptor proteins such BTB or SKP1, which, in turn, may recruit the Cullin machinery.

Activation of NF-κB signaling plays a major role in antiviral defenses as it controls the expression numerous effectors in the host antiviral defense system (25, 26). Consequently, many viruses have evolved strategies to manipulate key components of the NF-κB signaling pathway to evade host antiviral defenses and leverage it to enhance viral replication. One common strategy to disrupt NF-κB signaling is by inactivating upstream molecules of NF-κB signaling such as RLR and MAVS. For instance, HCV uses NS3/4a as protease to cleave MAVS. Another strategy is to prevent degradation of IκBα (26). For NF-κB to translocate into the nucleus, degradation of IκBα is required to release NF-κB. Rotaviruses inhibit NF-κB signaling by NSP1 protein mediated degradation of β-TrCP, an E3 ligase which is responsible to degrade IκBα (27, 28). Other viruses such as vaccinia virus use their A49 protein for molecular mimicry to sequester β-TrCP and stabilize IκBα (29). Additional strategies include direct targeting of NF-κB subunits, as exemplified by HSV-1 ICP0-mediated degradation of p50 or cytoplasmic sequestration of p65 by ectromelia virus EVM150 (30, 31). Viruses have also evolved mechanisms to disrupt NF-κB signaling at the level of the IKK complex. Hepatitis A virus 3C protease cleaves NEMO, and porcine reproductive and respiratory syndrome virus NSP4 binds NEMO to inhibit IKK activation(32, 33). Our findings identify reovirus σ3 as a distinct NF-κB signaling antagonist that targets this critical signaling node through a previously uncharacterized mechanism. Specifically, T3A σ3 associates with IKKβ and induces its degradation in a Cullin-RING ligase (CRL)-dependent manner, thereby suppressing NF-κB signaling.

As a viral protein, σ3 is multifunctional, reflecting the evolutionary pressure on viruses to encode many activities within limited genome space. Viral multifunctionality often emerges from flexible structural domains, modular interaction surfaces, and the ability to form distinct complexes. σ3 illustrates this well. It is one of the most abundant viral proteins, produced in many copies per infected cell. While most σ3 is incorporated into virions as a structural component required for particle stability and regulation of cell entry (13), a significant proportion remains available to modulate host responses. σ3 suppresses PKR activation and prevents stress granule formation, which has been linked to reovirus-induced myocardial injury (11). It also inhibits type I interferon signaling via strain-specific mechanisms: T3A σ3 degrades RIG-I and MAVS, whereas T3D^F^ and T1L σ3 inhibit transcriptional activity of NF-κB(10, 34, 35). Our discovery that T3A σ3 triggers CRL-dependent IKKβ degradation adds another layer to the multifunctional repertoire of this protein. Our work also highlights the remarkably different ways in which a highly homologous proteins from related reovirus strains inhibit the same pathway in distinct ways.

σ3 activity is further complicated by its ability to exist in several distinct configurations. It can remain as monomers, form homodimers, or assemble into a σ3-μ1 complex. The functional consequences of these forms differ substantially. For example, σ3 dimers bind dsRNA but σ3 cannot bind dsRNA when associated with μ1 (19). Consistent with this, we found that μ1 reduces the σ3-IKKβ interaction, suggesting that the free form, but not the σ3-μ1 complex targets IKKβ. σ3 and μ1 also differ in subcellular localization.

Some σ3 localizes diffusely in the cytoplasm, whereas σ3-μ1 complexes accumulate within lipid droplets or viral factories (17). Strain-specific differences in σ3 localization have been reported (36), raising the possibility that T3A σ3 remains more available in the free form to access host substrates such as the IKK complex. Although σ3 from both T3A and T3D^L^ strains can bind IKKβ, our qualitative analysis suggests a higher efficiency interaction by T3D σ3. Further, we find that only T3A σ3 induces IKKβ degradation. The two proteins differ by only nine amino acids, indicating that small polymorphisms may determine CRL recruitment, localization, or substrate recognition. Mapping σ3 residues required for IKKβ turnover will clarify how strain variation shapes immune evasion.

NF-κB signaling plays a critical role in determining disease outcomes during reovirus infection. In mice with neural cell-specific deletion of the NF-κB p65 subunit (Nsp65-/-mice, survival following intracranial reovirus infection is significantly increased compared to wild-type controls, indicating that p65 expression in neural cells enhances reovirus neurovirulence (37). This increased survival is accompanied by a marked reduction in virus-induced apoptosis in cortical neurons, despite comparable viral titers between wild-type and Nsp65-/-mice. This might be related to a reduction in the expression of genes involved in innate immunity, inflammation, and cell death in the neurons of Nsp65-/-mice (37). Studies in p50-null mice further highlight the tissue-specific roles of NF-κB signaling during reovirus infection. Similar to Nsp65-/-mice, p50 deficiency reduces apoptosis in the central nervous system, supporting a pro-apoptotic role for NF-κB in the brain (38). In contrast, loss of p50 in the heart leads to severe myocarditis, uncontrolled viral replication, and dramatically elevated cardiac viral titers than in wild-type mice, indicating that NF-κB p50 mediates a protective, pro-survival function in myocardial tissue. Notably, p50 deficiency has little effect on viral replication in the intestine or dissemination to secondary sites, emphasizing the organ-specific functions of NF-κB during reovirus infection (38). Given the importance of NFκB to reovirus pathogenesis, we speculate that viruses that differ in the capacity to block NFκB will display altered virulence. Such studies are a subject of our future investigation. We expect that findings from such studies will add to the growing evidence for σ3 as a virulence determinant (11, 39).

## MATERIALS AND METHODS

### Cells

L929 cells obtained from ATCC were maintained in Eagle’s MEM (Sigma-Aldrich) supplemented to contain 5% fetal bovine serum (FBS) (Life Technologies), and 2 mM L-glutamine (Invitrogen). HEK293T cells obtained from John Patton’s lab at Indiana University were maintained in Dulbecco’s MEM (Thermo-Fisher) supplemented to 10% FBS (Sigma-Aldrich, and 2mM L-glutamine (Invitrogen). ATCC L929 and HEK293T were used for all experiments to assess cell signaling. Spinner-adapted Murine L929 cells were maintained in Joklik’s MEM (Lonza) supplemented to contain 5% fetal bovine serum (FBS) (Sigma-Aldrich), 2 mM L-glutamine (Invitrogen), 100 U/ml penicillin (Invitrogen), 100 μg/ml streptomycin (Invitrogen), and 25 ng/ml amphotericin B (Sigma-Aldrich). Spinner-adapted L929 cells were used for cultivating, purifying and titering viruses.

### Plasmids

The empty vector pCMV-tag was obtained from Stratagene. pCMV-T3D^L^ S4 encoding for T3D^-L^ σ3 was generated through QuickChange mutagenesis. Plasmids for generating the T3D^L^ reverse genetics system were obtained from Takeshi Kobayashi (37). The T3A S4 gene encoding T3A σ3 was amplified from the TA-T3A-S4 plasmid and cloned into both the pT7 and pCMV vectors using In-Fusion Cloning. The pCMV-T3A S4 construct used for expression studies was similarly generated by In-Fusion Cloning. pT7-T3A S4 plasmid were used to generate the T3D^L^/T3A S4 monoreassortant virus by swapping the S4 gene of T3D^L^ S4 with pT7-T3A S4. pCI-T3D^F^ M2 encoding μ1 protein was obtained from Dr. John Parker’s lab. Expression vector for IKKβ-flag was purchased from Addgene (23298).

### Antibody and Reagents

Polyclonal antisera raised against T3D^L^ that have been described in the previous paper were used to detect viral proteins in infected cells(12). 4F2 monoclonal antibody to detect different strains of σ3 was obtained from University of Iowa Hybridoma Bank. Rabbit antisera specific for IKKβ were purchased from Cell Signaling Technology (Catalog no. 8943S). M2 Flag-tag antibody was purchased from Sigma-Aldrich (Catalog no. F3165). β-Actin antibody was purchased from ABclonal (Catalog no. AC026). MG132 and MLN4924 were purchased from Cayman Chemical Company and Sigma Aldrich and were used at a concentration of 25 μM and 2 μM. Cycloheximide was purchased from EMD-Millipore and was used at a concentration of 10 μg/mL. IKK inhibitor BAY-65-1942 (Bayer) was used at a final concentration of 10 μM. Custom synthesized siRNAs were purchased from Dharmacon. siRNA targeting β-galactosidase was used as control. The siRNA sequences used are as follows: β-galactosidase – CUACACAAAUCAGCGAUUU, T3D^L^ σ3 - CCUUAAACCUGAUGAUCGAUU, T3A σ3-CCUUAAACCUGAUGAUCGAUU. IR dye conjugated anti-mouse IgG and anti-rabbit IgG secondary antibodies were purchased from LICOR.

### Virus Purification

Prototype reovirus strains of T3D^L^ and T3D^L^/T3A S4 were regenerated by plasmid based reverse genetics (38) and a laboratory stock of T3A was obtained from T. Dermody’s laboratory. Purified reovirus virions were generated using second- or third-passage L-cell lysates stocks of reovirus. Viral particles were Vertrel-XF (Dupont) extracted from infected cell lysates, layered onto 1.2-to 1.4-g/cm^3^ CsCl gradients, and centrifuged at 187,183 x *g* for 4 h. Bands corresponding to virions (1.36 g/cm^3^) were collected and dialyzed in virion-storage buffer (150 mM NaCl, 15 mM MgCl_2_, 10 mM Tris-HCl [pH 7.4]) (39). Viral titer was determined by plaque assay using spinner-adapted L929 cells with chymotrypsin in the agar overlay.

### Growth Curve

L929 cells were seeded in 24-well plates (Greiner) at 2 × 10^5^ cells/well. 24 h after seeding, cells were adsorbed with virions T3D^L^ and T3D^L^/T3A S4 at an MOI of 1 PFU/cell for 1 h at 4°C. The cells were either lysed immediately after adsorption (input) or replaced with media as described above and incubated at 37°C (start of infection) for 24 or 48 h in a CO_2_ incubator. Afterwards, cells were harvested by 3× freeze-thaw cycle, and viral titters were measured by plaque assay on spinner cells. Briefly, confluent six-well plates of spinner cells were adsorbed with 100 µL of 10-fold serially diluted, freeze-thawed lysate for 1 h. Cells were then overlaid with Media 199 supplemented with 2 mM L-glutamine (Invitrogen), 0.5 U/mL penicillin, 50 µg/mL streptomycin (Sigma Aldrich), and 25 ng/mL amphotericin B (Sigma-Aldrich), and 1% Difco Bacto Agar (BD). Plates were incubated until countable plaques were visible and fixed with 3.7% formaldehyde in PBS. Fixed plates were then stained with 1% crystal violet in 5% ethanol solution to visualize and count plaques. The viral yield at a given time postinfection (*t*) was calculated using the following formula: log_10_(PFU/ml)*_t_*−log_10_(PFU/ml)_input_.

### ISVP Generation

ISVPs were generated in vitro by incubation of 2 x 10^12^ virions with 200 µg/ml of TLCK-treated CHT in a total volume of 0.1 ml at 32°C in virion storage buffer (150 mM NaCl, 15 mM MgCl_2_, 10 mM Tris-HCl [pH 7.4] for 30 min. Proteolysis was terminated by addition of 2 mM phenylmethylsulphonyl fluoride (PMSF) and incubation of reactions on ice. Generation of ISVPs was confirmed by SDS-polyacrylamide gel electrophoresis (PAGE) and Coomassie Brilliant Blue staining.

### Plasmids and siRNA Transfection

In 12-well plates, HEK293T were seeded in the previous day to reach 80%-90% confluent at the time of transfection. Afterwards, the serum media was replaced by OPTI-MEM prior to transfection. 4 μL of Lipofectamine 2000 (Thermo-Fishers) was used to transfect various plasmids with total concentration up to 1.6 ug according to manufacturer instructions for different plate sizes. After 25 minutes of incubation at room temperature, the transfection complexes were overlayed on the top of cells and incubated for 5 h. The OPTI-MEM media was replaced by serum media after 5hpt and incubated at 37°C for 24-48h to obtain optimum protein expression. When needed, the transfected cells were treated with MG132, MLN4924, or DMSO within indicated time intervals. For siRNA transfection, in 6-wells plates, 8μL of INTERFERin (Polyplus) was used to transfect siRNA to final concentration of 40nM according to the manufacturer instructions. 40% of cells (5x10^5^ cells/mL) were seeded on top of the siRNA-lipofectamine/INTERFERin mixture. Virus infection was performed 28-48 h following siRNA transfection.

### Infection and Preparation of Cellular Extracts

Cells were either adsorbed with PBS or reovirus at room temperature for 1 h, followed by incubation with media at 37°C for the indicated time interval. When needed, DMSO or MLN4924 were added to the media immediately after the 1 h adsorption period and incubated for 20 h post infection. After viral adsorption, the cycloheximide or DMSO was added and incubated for 20 h. For preparation of whole cell lysates, cells were washed in phosphate-buffered saline (PBS) and lysed with 1X RIPA (50 mM Tris [pH 7.5], 50 mM NaCl, 1% TX-100, 1% DOC, 0.1% SDS, and 1 mM EDTA) containing a protease inhibitor cocktail (Roche) and 2 mM PMSF, followed by centrifugation at 15000 × *g* for 10 min at 4°C to remove debris. The lysates will then be used for immunoblotting and immunoprecipitation.

### Immunoprecipitation Assay

The cell was lysed with 1X RIPA buffer and protease cocktail inhibitor (Roche). Cells lysates were immunoprecipitated with α-flag M2 antibody (Sigma Aldrich) or 4F2 σ3 monoclonal antibody with concentration of 2ug and incubated overnight at 4°C in the rotator. Protein A/G magnetic beads (Pierce) were pre-washed and incubated with antibody-lysate for 2 h. Afterwards, the beads-lysate mixtures were washed with wash buffer 3 times for 5 min. The beads-immunoprecipitated mixtures were incubated in elution buffer for 20 minutes at 70°C and eluted to obtain immunoprecipitated lysates.

### Immunoblot Assay

Cell lysates were resolved by electrophoresis on 10% polyacrylamide gels and transferred to nitrocellulose membranes. Membranes were blocked for at least 1 h in TBS containing T20 Starting Block (Thermo Fisher) or 2% BSA and incubated with antisera against reovirus (1:5000), σ3 (1:1000), IKKβ(1:1000), Flag-tag (1:1000), or β-actin (1:10000) at 4°C overnight. Membranes were washed three times for 10 min each with washing buffer (TBS containing 0.1% Tween-20) and incubated with 1:10000 dilution of Alexa Fluor conjugated goat anti-rabbit IgG or goat anti-mouse IgG depending on the primary antibody in blocking buffer. Following three washes, membranes were scanned using an Odyssey Infrared Imager (LI-COR). Band intensity was analyzed using Image Studio Lite software (LICOR).

### Immunofluorescence Microscopy

L929 cells were grown on fibronectin pretreated eight-chamber cover glasses (Nunc Lab-Tech, ThermoFisher Scientific) and infected with Reovirus T3D^L^/T3A S4 at MOI 10. After adsorption, the cells were washed with PBS and treated with either DMSO or MLN4924. After 24 h, cells were washed with PBS and treated with 4% paraformaldehyde. Then, the cells were blocked with 5% normal donkey serum with 0.05% Triton X100 in PBS (Blocking buffer) followed by permeabilization with 0.1% saponin and 0.1% Triton X100. Primary antibodies were diluted in the Blocking buffer and incubated the cells for 1h in room temperature. After three washes, cells were labeled with secondary antibody for 1h in room temperature. Finally, the nucleuses were stained with Hoechst 33342 (Tocris Bioscience) and washed with PBS. Samples were imaged using Leica Stellaris 8 confocal microscopy running by LASX software. Then the images were imported to a Huygens Essential v17.10 software and deconvolved (Scientific Volume Imaging). The deconvolved images were imported to ImageJ software to add scale bars.

### RT-qPCR

RNA was extracted from infected cells at various time intervals after infection using Total RNA mini kit (Biorad). For RT-qPCR, 0.5 to 2 µg of RNA was reverse transcribed with the High-Capacity cDNA Reverse Transcription Kit (Applied Biosystems) using random hexamers for amplification of cellular genes. A 1:10 dilution of the cDNA was subjected to PCR using SYBR Select Master Mix (Applied Biosystems). Fold increase in gene expression with respect to control sample (indicated in each figure legend) was measured using the ΔΔC_T_ method(40).

### Statistical Analysis

Statistical significance between experimental groups was determined using the unpaired student’s *t*-test, one-way ANOVA, or two-way ANOVA function and graphed using Graphpad Prism software. Statistical analyses for differences in gene expression by RT-qPCR were done on the ΔΔC_T_ values.

